# Mevastatin in colon cancer by spectroscopic and microscopic methods - Raman imaging and AFM studies

**DOI:** 10.1101/2021.08.31.458425

**Authors:** K. Beton, P. Wysocki, B. Brozek-Pluska

## Abstract

One of the most important areas of medical science is oncology, which is responsible for both the diagnostics and treatment of cancer diseases. Simultaneously one of the main challenges of oncology is the development of modern drugs effective in the fight against cancer. Statins are a group of biologically active compounds with the activity of 3-hydroxy-3-methyl glutaryl-CoA reductase inhibitors, an enzyme catalyzing the reduction of 3-hydroxy-3-methyl-glutaryl-CoA (HMG-CoA) to mevalonic acid. By acting on this enzyme, statins inhibit the endogenous cholesterol synthesis which in turn causes the reduction of its systemic concentrations. However, in vitro and in vivo studies confirm also the cytostatic and cytotoxic effects of statins against various types of cancer cells including colon cancer. In the presented studies the influence of mevastatin on cancerous colon cells CaCo-2 by Raman spectroscopy and imaging is discussed and compared with biochemistry characteristic for normal colon cells CCD-18Co. Based on vibrational features of colon cells: normal cells CCD-18Co, cancerous cells CaCo-2 and cancerous cells CaCo-2 treated by mevastatin in different concentrations and incubation times we have confirm the influence of this statin on biochemistry composition of cancerous human colon cells. Moreover, the spectroscopic results for colon normal cells and cancerous cells based on data typical for nucleic acids, proteins, lipids have been compared. The cytotoxisity of mevastatin was determined by using XTT tests.

## Introduction

Colorectal cancer (CRC) is one of the most frequently detected neoplastic changes. It is also one of the most difficult types of cancer to cure, as evidenced by the very large number of deaths caused by this disease [1–4]. CRC is formed based on the benign adenomas - polyps [1]. The tumor formation most often is the result of genetic transformation of normal cells. The genes involved in mutations that lead to cancer include oncogenes, suppressor genes (anti-oncogenes), and mutator genes (DNA repair genes). As the result of this mutation the conversion of an initially normal cells into a cancerous one, and the subsequent formation of metastases can be observed.

The basic method of oncological treatment of the colon cancer is resection, i.e. removal of a part covered by the tumor. To extend the patients survival rate radiation therapy also can be applied, reducing the frequency of cancer relapses. In addition, chemotherapy can be included. The most commonly substances used in the treatment of colon cancer are: fluorouracil (FU) and calcium folinate (FA). They are used in a standard chemotherapy procedure. In order to increase the effectiveness, a method of multi-drug therapy also has been developed. In addition to FU / FA treatment irinotecan and oxaliplatin can be used. Thanks to the use of these compounds the survival rate of patients affected by cancerous changes in the colon may be increased.

Statins occur both naturally and are synthesized in laboratories. Depending on the structure of the statin compound, mevastatin, lovastatin, pravastatin, compactin (statins of natural origin), simvastatin (semi-synthetic statins), atorvastatin, rosuvastatin, pitavastatin, cerivastatin and fluvastatin (synthetic statins) can be distinguished [5]. All these compounds have a pharmacophore group in their active form. Most often, statins of natural origin come from all kinds of fungi, especially mold fungi [6]. In their case, the structural basis is partially hydrogenated naphthalene having a hydroxyl group, which is esterified with a 2,2-dimethylbutyric acid residue or a 2-methylbutyric acid residue in the 1-position, a pharmacophore group (a six-membered lactone ring having a hydroxyl group - in the case of pravastatin and in active forms mevastatin, simvastatin and lovastatin, the ring is hydrolyzed to the β-hydroxy carboxylic acid form at position 4 and with a methyl group at position 7). For synthetic statins, the basic building unit of the molecule is different for each type of compound. For fluvastatin it is indole, for rosuvastatin it is pyrimidine, for atorvastatin it is pyrrole, for cerivastatin it is pyridine and for pitavastatin it is quinoline [7]. Each of these compounds is substituted with a pharmacophore group (identical to the structure of natural statins) and a p-fluorophenyl group. It can therefore be concluded that the differences between synthetic statins are much more noticeable than with natural statins. The structure of the molecule of each statin is very similar to 3-hydroxy-3-methylglutaryl coenzyme A (3-HMG-CoA).

In addition, statins show a very high affinity for 3-hydroxy-3-methylglutaryl coenzyme reductase, which is why they bind to it through a reversible reaction as an inhibitor. By acting on this enzyme, statins inhibit the endogenous cholesterol synthesis which in turn causes the reduction of its systemic concentrations. However, statins show not only cholesterol modulating properties, their show even more valuable features as anti-cancer drugs caused by their influence on proteins of the Ras superfamily, which are small monomeric messenger proteins located on the cytoplasmic side of the cell membrane. They have the ability to attach and hydrolyze GTP [8–14]. By activating MAPK signaling pathways (Ras / Raf / MEK / ERK1 / 2, PI3K / Akt, JAK / STAT etc.) proteins influence the processes related to the growth, differentiation and apoptosis of eukaryotic cells. On the other hand, Rho proteins, mediating the activation of the kinase FAK (adhesion contact kinase), ROCK (Rho-dependent protein kinase) and PAK kinases (P21ras-activated kinase), modulate primarily the actin cytoskeleton [10,15–17]. Thus, the Rho protein influences the functioning of the contractile system in cells muscles and the cytokinetic spindle in division cells. As second order messengers, they participate in the regulation of vesicular transport and the cell cycle, and mediate cell migration and adhesion to extracellular matrix components. Taking into account the variety of GTPase-activated Ras and Rho-enzyme pathways, it seems obvious that their mutations can be the cause of numerous pathological processes in the body. Frequent occurrence mutated Ras or Rho in various types of cancer indicates the possibility of using statins in inhibition the pro-oncogenic properties of these proteins and their prevention in tumor development [18].

Statins can be used as anti-cancer compounds also thanks to their ability to inhibit the multiplication (proliferation) of neoplastic cells. In the work by A. Sławińska et al. [19] experiments on cells derived from rats (HTC-4 - liver cancer), mice (LLC-L1 - lung cancer; MCA-38 - colorectal cancer, B16F10 - melanoma) have been presented. The cells were treated with lovastatin. Depending on the dose of the statin compound, inhibition of cell growth and a decrease in their viability was observed. Lovastatin showed antiproliferative activity at low doses, and pro-apoptotic activity at higher doses. In addition to animal cell studies, studies on cells of human origin: the leukemia line (HL-60, THP-1, Jurkat), myeloma line (U266, RPMI8226, MCC-2), breast cancer (MDA-MB-237, MCF-7), colon (CaCo-2) , bladder (T24) and prostate (DU-145, PC-3) have been carried out. The results performed with human cells were consistent with those obtained with cells of animal origin. The high cytotoxicity of statins towards neoplastic cells may be a consequence of both the disturbance of the cholesterol production process and the inhibition of the production of isoprenoid compounds by the mevalon pathway [19]. Another feature presented in the work of Sławińska et al. is the effect of statin compounds on DNA synthesis. Based on studies in which lovastatin was applied to human cervical cancer (HeLa) cells and human breast adenoma (MCF-7) cells, consistent results of a 90-98% decrease in DNA synthesis was observed. However, there was a difference in the results when studies were carried out on malignant cells isolated from the blood of patients suffering from different types of myeloid leukemia. The differences were manifested in a reduced level of incorporated 3H thymidine in the majority of the tested cells (about 60%). As a result of the disturbance in the synthesis of isoprenoid compounds, statins made it impossible to farnesylate prelines, i.e. filament proteins derived from intermediate nuclear matrices, and to bind them to the nuclear envelope. As a result, most likely there was a disintegration of the cell nucleus and, consequently, a large number of changes in both the chromatin structure and gene expression. Another explanation for this phenomenon may be the blockage of the cell cycle in the G2 phase. In turn, as a result of studies of mouse lung cancer cells (LLC-L1) and human liver cancer cells (HTC-4), an increase in the percentage of cells labeled with thymidine was observed depending on the increase in the dose of lovastatin. The differences in the investigated level of cells labeled with thymidine can be explained by the stimulating effect of statin compounds on the activity of processes responsible for the regeneration of genetic material. It has also been proven in numerous studies that the action of lovastatin on cells for 24 hours suppresses cell growth in the G1 / S phase, and exposure of cells to the action of lovastatin for 5 days resulted in the growth of cells blocked in the G2 phase. As a result of this, the number of mitotic cells was reduced [20–22].

Another feature of statin compounds that makes them useful in cancer therapy is their ability to induce apoptosis, that is, the ability to control the removal of used or damaged cells from a multicellular organism by killing them to heal the body. A very large number of studies have clearly demonstrated the pro-apoptotic effect of statins on cancer cells of the larynx, pancreas, cervix, colon, thyroid, prostate, malignant glioma and breast [16–22]. Moreover, the apoptotic effect was also observed in the case of cell cultures derived from acute myeloid leukemias, lymphoblastic leukemias and multiple myeloma cells. An important observation of great importance for cancer therapy is that statins have an apoptotic effect mainly in the case of neoplastic cells. This means that statins are not destructive in normal cells, as is the case with most drugs used to treat cancer [23–25].

During the course of neoplastic processes, significant changes take place in the expression of growth factors, surface receptors, adhesion molecules, cytokines or proteliotic enzymes, as a result of which the cell acquires an invasive phenotype that enables the metastasis of neoplastic cells. An important feature of malignant (metastatic) cells is the loss of adhesion properties and the acquisition of the ability to create capillaries (angiogenesis), which also results in the formation of blood vessels that are needed for tumor growth. Another important feature of malignant neoplastic cells is the secretion of metalloproteinases, enzymes involved in the degradation and rebuilding of the extracellular matrix and the basement membrane (collagen, fibronectin, elastin, laminin, proteoglycans and glycoproteins). These enzymes are created by other types of cells, including epithelial cells, fibroblasts, endothelial cells, myocytes, neurons, cells of the immune system (lymphocytes, monocytes, dendritic cells, granulocytes) [19].

Due to pleiotropic properties and low cytotoxicity to cells with normal structure, statin compounds constitute a very attractive group of compounds that may become common drugs inhibiting invasive neoplasms. On the basis of molecular studies using the methods of zymography, immunoblotting and real-time polymerase chain reaction (RT-PCR), it was proved that statin compounds inhibited the expression and activity of many metalloproteinases. It has also been found to inhibit the invasiveness of the malignant phenotype of neoplastic cells from various sources after treatment with lovastatin, for example in a patient with leukemia, glioblastoma or malignant melanoma. A similar relationship was observed after the action of fluvastatin and cerivastatin on pancreatic, breast and colorectal cancer cells [26–28].

One of the main reasons for the anti-tumor activity of statin compounds is their counteracting the formation of new blood vessels in the tumor area. In other words, statins inhibit angiogenesis. In turn, in tests on colon cancer cells, a decrease in the secretion of platelet PDGF growth factor was observed. Studies in which statins were combined with bisphosphonate led to a reduction in the expression of the factor responsible for the growth of fibroblasts and integrin, which resulted in a reduction in the ability of cancer cells to metastasize [18,21,29,30].

As a result of research on cancer cells of various types, the influence of statin compounds on changes in the organization of the cytoskeleton of cells was also observed. The result was the destabilization of actin filaments and the reduction of cytoplasmic protrusions as a result of the disappearance of stress fibers. In addition, a change in cell morphology, loss of mobility and the acquisition of features characteristic of apoptotic cells were also observed. Another very interesting feature observed in the research of melanoma, brain, liver and lung cancers is the ability of statin compounds to stop metastases of invasive types of solid tumors. Significant reductions in tumor mass and an increase in patients’ lifespan have also been observed [31]. However, much more research is needed to understand each of the statin compounds to the highest degree, allowing them to be safely used in cancer therapy.

In this paper the influence of mevastatin on cancerous colon cells CaCo-2 by Raman spectroscopy and imaging are discussed and compared with biochemistry typical for normal colon cells CCD-18Co. Based on vibrational features of colon cells: normal cells CCD-18Co, cancerous cells CaCo-2 and cancerous cells CaCo-2 treated by mevastatin in different concentrations and incubation times we have confirm the influence of this statin on biochemistry composition of cancerous cells. Moreover, the spectroscopic results for colon normal cells and cancerous cells based on data typical for nucleic acids, proteins and lipids have been compared. The cytotoxisity of mevastatin was determined by using XTT tests. Additionally, using AFM technique we have analyzed nanomechanical properties of CCD-18Co, CaCo-2 cells including CaCo-2 supplemented by mevastatin. The topography, Young’s modulus and adhesion for all normal and cancer human colon cells were also recorded and analyzed.

## Materials and methods

### Cell lines and cell culture

CCD-18Co cell line (ATCC® CRL-1459™) was purchased from ATCC: The Global Bioresourece Center. CCD-18Co cell line was cultured using ATCC-formulated Eagle’s Minimum Essential Medium with L-glutamine (catalog No. 30-2003). To make the complete growth medium, fetal bovine serum was added to a final concentration of 10%. Every 2–3 days, a new medium was used. The cells obtained from the patient are normal myofibroblasts in the colon. The biological safety of the CCD-18Co cell line has been classified by the American Biosafety Association (ABSA) as level 1 (BSL-1). The CaCo-2 cell line was also purchased from ATCC and cultured according to the ATCC protocols. The CaCo-2 cell line was obtained from a patient - a 72-year-old Caucasian male diag-nosed with colon adenocarcinoma. The biological safety of the obtained material is clas-sified as level 1 (BSL - 1). To make the medium complete we based on Eagle’s Minimum Essential Medium with L-glutamine, with addition of a fetal bovine serum to a final concentration of 20%. The medium was renewed once or twice a week.

### Cultivation conditions

Cell lines (CCD-18Co, Caco-2) used in the experiments in this study were grown in flat-bottom culture flasks made of polystyrene with a cell growth surface of 75 cm^2^. Flasks containing cells were stored in an incubator providing environmental conditions at 37 °C, 5% CO_2_, 95% air.

### Raman Spectroscopy and Imaging

All maps and Raman spectra presented and discussed in this paper were recorded using the confocal microscope Alpha 300 RSA+ (WITec, Ulm, Germany) equipped with an Olympus microscope integrated with a fiber with 50 μm core diameter with a UHTS spectrometer (Ultra High Through Spectrometer) and a CCD Andor Newton DU970NUVB-353 camera operating in default mode at −60 °C in full vertical binning mode. 532 nm excitation laser line, which is the second harmonic of the Nd: YAG laser, was focused on the sample through a Nikon objective lens with magnification of 40x and a numerical aperture (NA = 1.0) intended for cell measurements performed by immersion in PBS. The average excitation power of the laser during the experiments was 10 mW, with an integration time of 0.5 s for Raman measurements for the high frequency region and 1.0 s for the low frequency region. An edge filter was used to filter out the Rayleigh scattered light. A piezoelectric table was applied to set the test sample in the right place by manipulating the XYZ positions and consequently record Raman images. Spectra were acquired with one acquisition per pixel and a diffraction grating of 1200 lines/mm. Cosmic rays were removed from each Raman spectrum (model: filter size: 2, dynamic factor: 10) and the Savitzky-Golay method was implemented for the smoothing procedure (order: 4, derivative: 0). All data was collected and processed using a special original software WITec Project Plus. All imaging data were analyzed by Cluster Analysis (CA), which allows for grouping of a set of vibrational spectra that bear resemblance to each other. CA was executed using WITec Project Plus software with Centroid model and k-means algorithm, in which each cluster is represented by one vector of the mean. Data normalization was performed using the model: divided by norm, which was wrought with the Origin software serving as mathematical and statistical analysis tool. The Origin software was also used to perform ANOVA analysis necessary to indicate statistically significant results (means comparison: Tukey model, significance level: 0.05).

### AFM measurements

AFM measurements were performed by using PIK Instruments atomic force microscope with scanning range of 100 x 100 μm in the X and Y axes and 15 μm in the Z axis with a positioning resolution in the XY axis of 6 pm and in the Z axis of 0.9 pm, equipped with an inverted microscope, enabling measurements in air and liquid, in contact and tapping modes. Nanosurf C3000 software was used for AFM data collection. During measurements topography maps and nanomecanical properties of cells with and without supplementation of mevastatin have been determined with the resolution 256×256 point per 60×60 m. During measurements qp-Bio-AC-50 tips produced by Nanosensors with a spring constant 0.6 N/m were used. The analysis of AFM data was performed by using AtomicJ software [32] to obtain information about Young’s modulus and adhesion of analysed biological samples.

For AFM measurements cells were cultured on Petri dishes filled with culture media. Once the growing cells formed semi-confluent monolayer, the dish with cells was mounted on the AFM scanner, medium was replaced by PBS and the sample was measured within next 2–3 h at room temperature and ambient conditions.

### Chemical compounds

Mevastatin ≥98% (HPLC), powder or crystals catalogue number M2537-5MG, bisBenzimide H 33342 trihydrochloride catalogue Number B2261, Red Oil-O catalogue Number O0625 were purchased from Merck Life Science Sp. z o. o., and used without additional purification. XTT proliferation Kit with catalogue Number 20-300-1000 was purchased from Biological Industries.

### XTT

In order to be able to perform clinical trials, a series of tests should be carried out to determine the activity of cells in terms of their metabolism and proliferation after exposure to specific substances [33]. This is necessary because on this basis it is possible to determine whether a given chemical is producing a cytotoxic response. Initially, tests were developed to incorporate compounds such as 5-bromo-2-deoxyuridine (BrdU) or (H) -thymidine into the structure of DNA [34]. Due to the inconvenience of this type of tests related to the need to use radioactive materials, expensive equipment or a time-consuming procedure, colorimetric methods have been developed. The basis of this method is the phenomenon observed for tetrazolium salts, which can be transformed by living cells as electron acceptors. As a result of this transformation, colored formazan compounds are formed. The first salt to be used in the colorimetric tests is 3-(4,5-dimethylthiazol-2-yl)-2,5-diphenyltetrazolium bromide known as the MTT salt. It is a positively charged compound, thanks to which it easily penetrates the cell, where it is reduced to a water-insoluble formazan compound [35]. However, this method is also not perfect due to the need to dissolve the formazan compound crystals in an organic solvent. For this reason, a method was developed in which the MTT salt was replaced with the 2,3-bis-(2-methoxy-4-nitro-5-sulfophenyl)-2H-tetrazoli-5-carboxanilide sodium salt, more widely known as the XTT salt. Unlike MTT, the XTT salt, when it enters the cell, is transformed into a product that can be dissolved in an aqueous medium. XTT, unlike the MTT salt, has a negative charge, so its permeability to the cell interior is low [36]. This results in a reduction either at the cell surface or in the plasma membrane by the transmembrane electron transport chain.

One application of the XTT colorimetric assay is to test the viability of cells as a function of the compound that is active on them and the concentration of the compound. An example of this type of compound is statins [37]. In the publication by Ludwig et al. the effect of three statins was investigated: atorvastatin, simvastatin and pravastatin [37]. For this purpose, a test was performed for each of the compounds for different concentrations of the test substance. The statin compound was added after placing normal endothelial cells (CPAE) in a 96-well plate medium and after incubating the cells for 24 h. In addition, a control was performed with only cells submerged in the medium. Then, after the addition of statins, the cells were incubated again for 4 h, after which it was possible to perform the measurement. Based on the obtained results, the survival curves of the studied cells were determined depending on the statin compound used.

### Determination of the appropriate mevastatin concentration using the XTT test

For each cell type, XTT tests were performed 24 h and 48 h after the addition of mevastatin to the cells immersed in the culture medium. Preparation for the test included proper filling of the 96-well plate according to the procedure developed at the Institute of Applied Radiation Chemistry in Lodz. The wells were filled in such a way that each row contained a specific series of measurements. For example - in one row all plates were filled with medium, in another - control samples containing only cells immersed in the medium, and only in subsequent rows - cells in the medium with the addition of a specific concentration of mevastatin. 6 different concentrations of mevastatin were selected for the test: 1 μM, 5 μM, 10 μM, 25 μM, 50 μM, 100 μM. After completing each of the 96-well plates, the samples were incubated at 37 °C. After the time from the addition of mevastatin (24 and 48 h), the XTT test was performed using the BioTek Synergy HTX apparatus. The measurement took about 3 hours. After the completion of the study, the obtained results had to be analyzed using a spreadsheet, resulting in a bar graph showing the effect of mevastatin concentration on the survival of the tested cell type, taking into account the time since the addition of mevastatin.

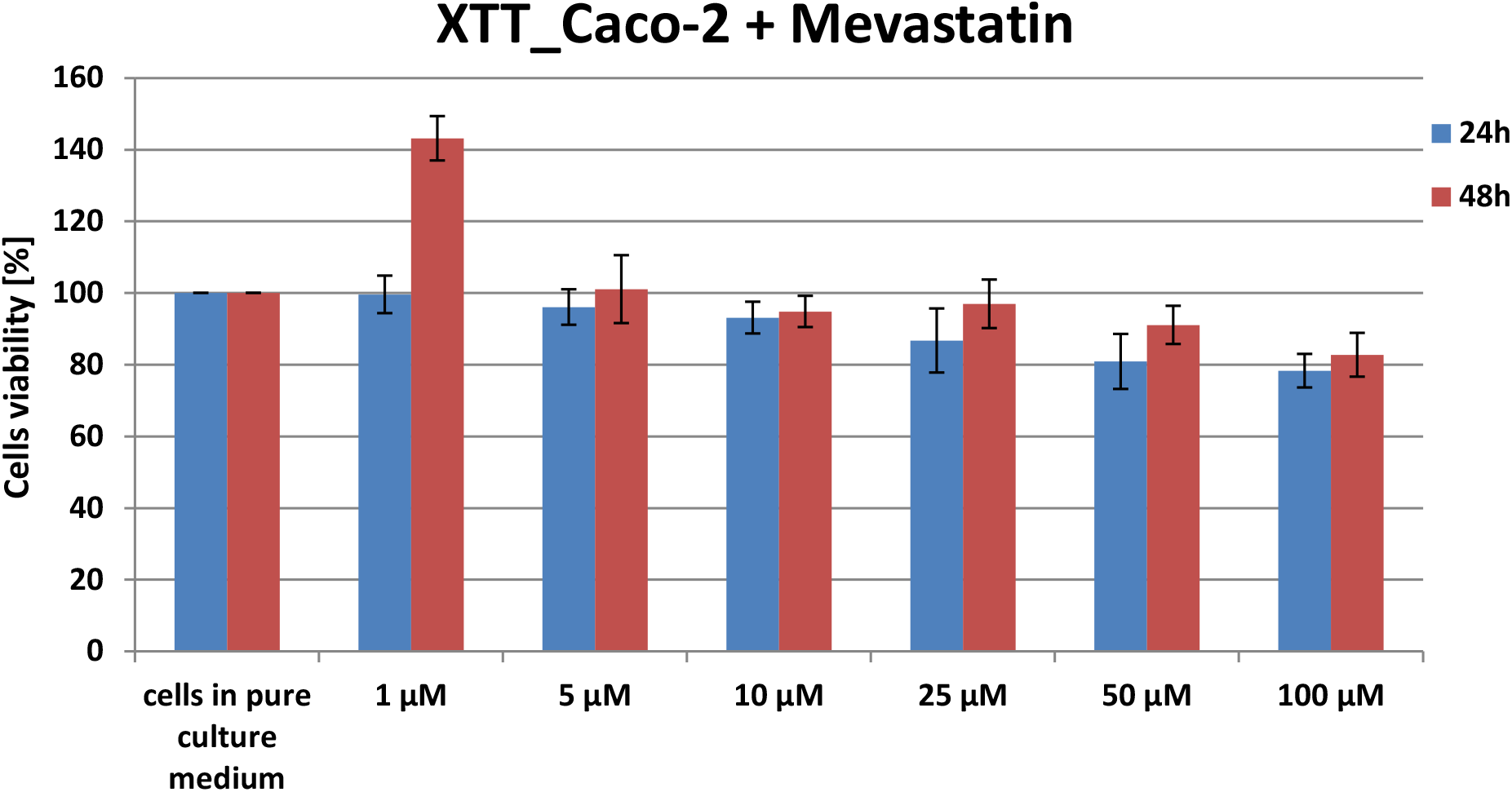

For both normal (CCD-18Co) and cancerous (CaCo-2) cells, it was found that the most appropriate concentration of mevastatin in the cell solution with medium would be 10 μM. For each test, cell survival at such a concentration fluctuated in the range of 50-60%, which made it possible to conclude that at such a concentration, the effect of mevastatin on cells will be noticeable both in the study of nanomechanical and biochemical properties, and there will be enough living cells to allow will be conducting analyzes. The exception were CCD-18Co cells 24 h after the addition of mevastatin - for this test, cell survival at a mevastatin concentration of 10 μM was approximately 37%. In this study, in the further part of the experiments (Raman spectroscopy; AFM), the effect of mevastatin only on colon cancer cells (CaCo-2) after 24 and 48 hours was investigated.

### Results and discussion

One of the main goals of our study was to determine the statistically significant differences between normal and cancer human colon cells including cancer cells supplemented by mevastatin based on vibrational features of them. Therefore to properly address these tasks we investigated systematically how the Raman method responds to in vitro normal and cancer human cells with and without the supplementation by mevastatin.

Figure 1 shows the microscopy image, Raman image, Raman images of all cells substructures identify by using Cluster analysis algorithm, average Raman spectra typical for identified: lipid rich structures, mitochondria, nucleus, cytoplasm and cell membrane, cell environmental, and the average Raman spectra for the cell as a whole for human normal colon cells: CCD-18Co (panel I) and human cancer colon cells CaCo-2 (panel II).

**Figure 1.**
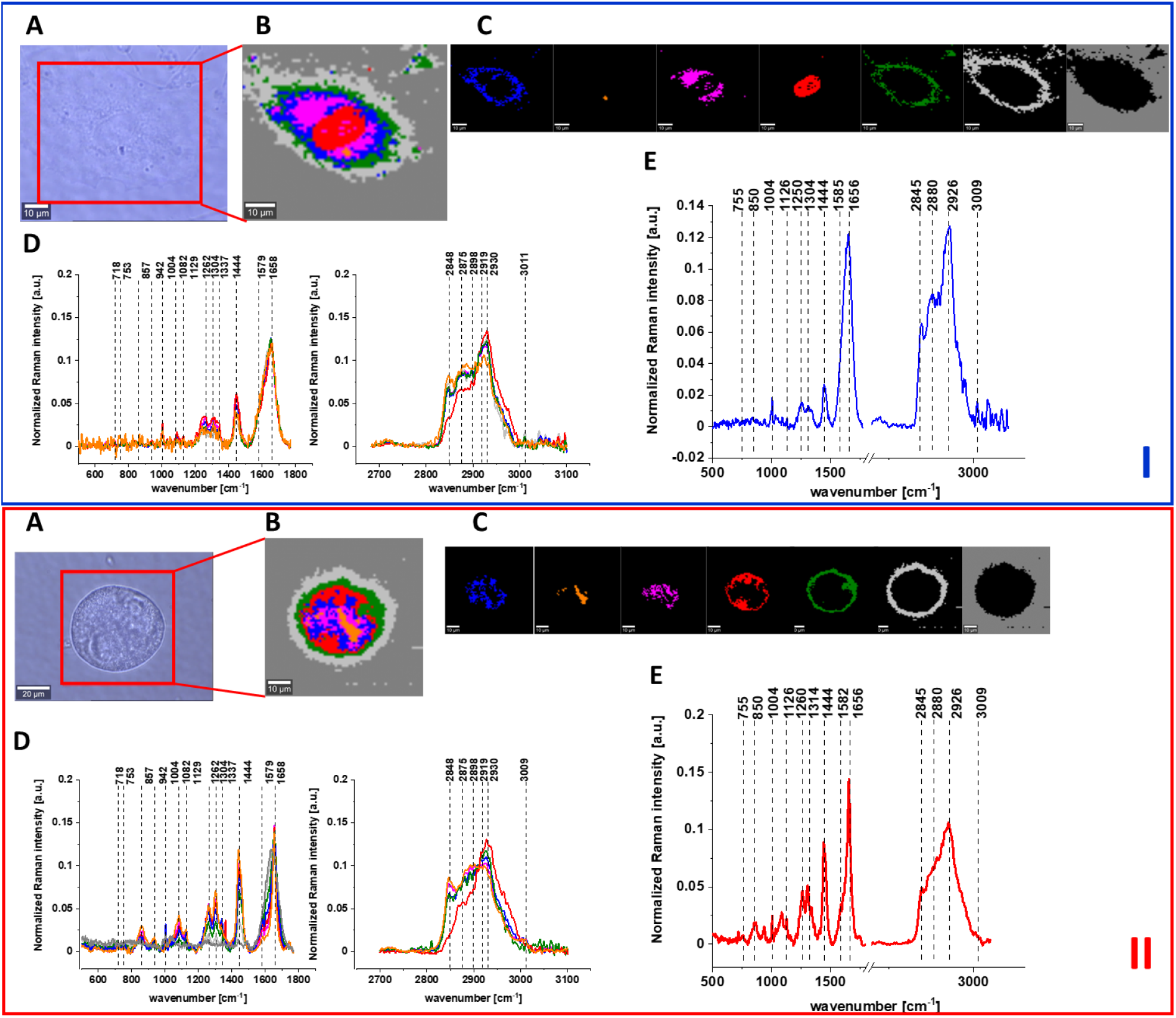
The microscopy image (A), Raman image (B) constructed based on Cluster Analysis (CA) method, Raman images of all clusters identified by CA assigned to: lipid-rich regions (blue and orange), mitochondria (magenta), nucleus (red), cytoplasm (green), cell membrane (light grey), and cell environment (dark grey) (C), the average Raman spectra typical for all identified clusters for low frequency and high frequency region (D), and the average Raman spectrum for cell as a whole (E) for human colon normal single cell CCD-18 Co (I) and cancer human colon single cell (II). All data were obtained for experiments without any supplementation, cells measured in PBS, colors of the spectra correspond to the colors of clusters, excitation laser line 532 nm, the average Raman spectra calculated based on the data for 6 cells.

One can see from Figure 1 that using Raman image mode the human colon normal CCD-18Co and human colon cancer cells CaCo-2 can be characterized by high resolved vibrational spectra.

The fingerprint region of Raman spectra provides basic information on the biochemical composition of analyzed sample e.g. the peak at 755 cm^−1^ is associated with nucleic acids, DNA, tryptophan, nucleoproteins [38,39], broad peak around 850 cm^−1^ is assigned to tyrosine [38], a sharp peak at 1004 cm^−1^ corresponds to the aromatic amino acid phenylalanine [39] while peak at 1126 cm^−1^ is typical for saturated fatty acids, band at 1304 cm^−1^ corresponds to CH_2_ deformation vibration of lipids, adenine, cytosine [39], band at 1444/1452 cm^−1^ is typical for lipids and proteins, peak at 1585 cm^−1^ for CN_2_ scissoring and NH_2_ rock of mitochondria and phosphorylated proteins [38,40,41]. Bands characterizing protein secondary structure are also observed for fingerprint region: the Amide I (C=O stretch) near 1656 cm^−1^, the Amide II (N–H bend + C–N stretch) near 1557 cm^−1^, and very weak the Amide III (C–N stretch + N–H bend) near 1260 cm^−1^ [39,42,43]. The high frequency peaks originates in the symmetric and antisymmetric stretching vibrations of C-H bonds found in lipids, glycogen, proteins, RNA, and DNA. Lipids and fatty acids including unsaturated fraction indicating the presence of fat appear at 2845, 2880, 3009 cm^−1^. Proteins contribution, in the high frequency region, is observed at 2875, 2888, 2919 and 2926 cm^−1^ [39,44].

The same Raman spectroscopy based techniques were applied to analysis of vibrational features of cancer human colon cells CaCo-2 after adding of mevastatin with concentration of 10 μM for 24 and 48h.

Figures 2 and 3 show the microscopy image, Raman image, Raman images of all cell substructures identified by using Cluster analysis algorithm, average Raman spectra typical for identified: lipid rich structures, mitochondria, nucleus, cytoplasm and cell membrane, cell environmental, the average Raman spectra for the cell as a whole of cancer human colon cells after adding of mevastatin in the concentration of 10 μM for 24 and 48h respectively.

**Figure 2.**
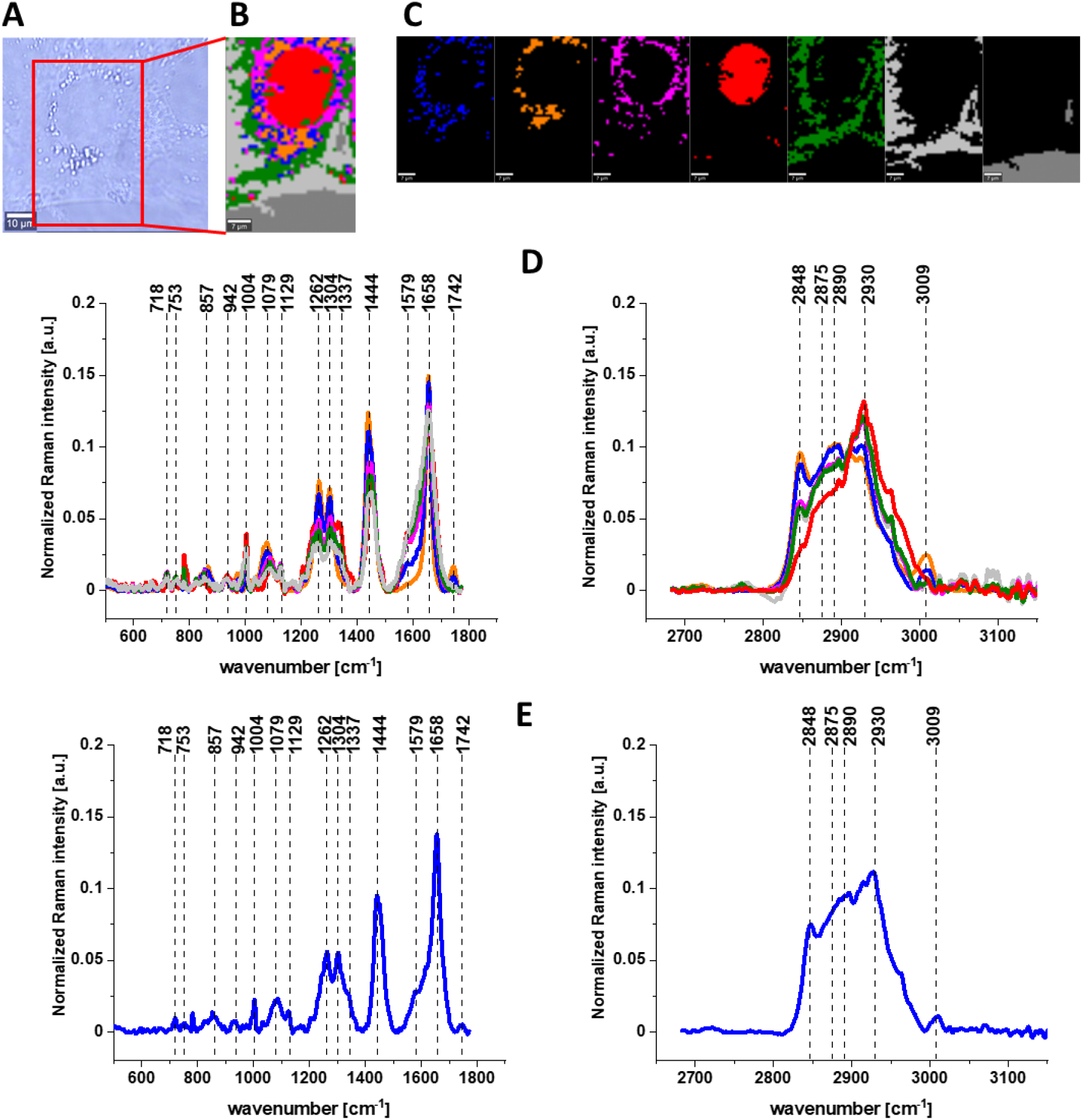
The microscopy image (A), Raman image (B) constructed based on Cluster Analysis (CA) method, Raman images of all clusters identified by CA assigned to: lipid-rich regions (blue and orange), mitochondria (magenta), nucleus (red), cytoplasm (green), cell membrane (light grey), and cell environment (dark grey) (C), the average Raman spectra typical for all identified clusters for low frequency and high frequency region (D), and the average Raman spectrum for cells as a whole (E) for cancer human colon cells after adding of mevastatin in 10 μM concentration for 24h. Cell measured in PBS, colors of the spectra correspond to the colors of clusters, excitation laser line 532 nm, the average Raman spectra calculated based on the data for 6 cells.

**Figure 3.**
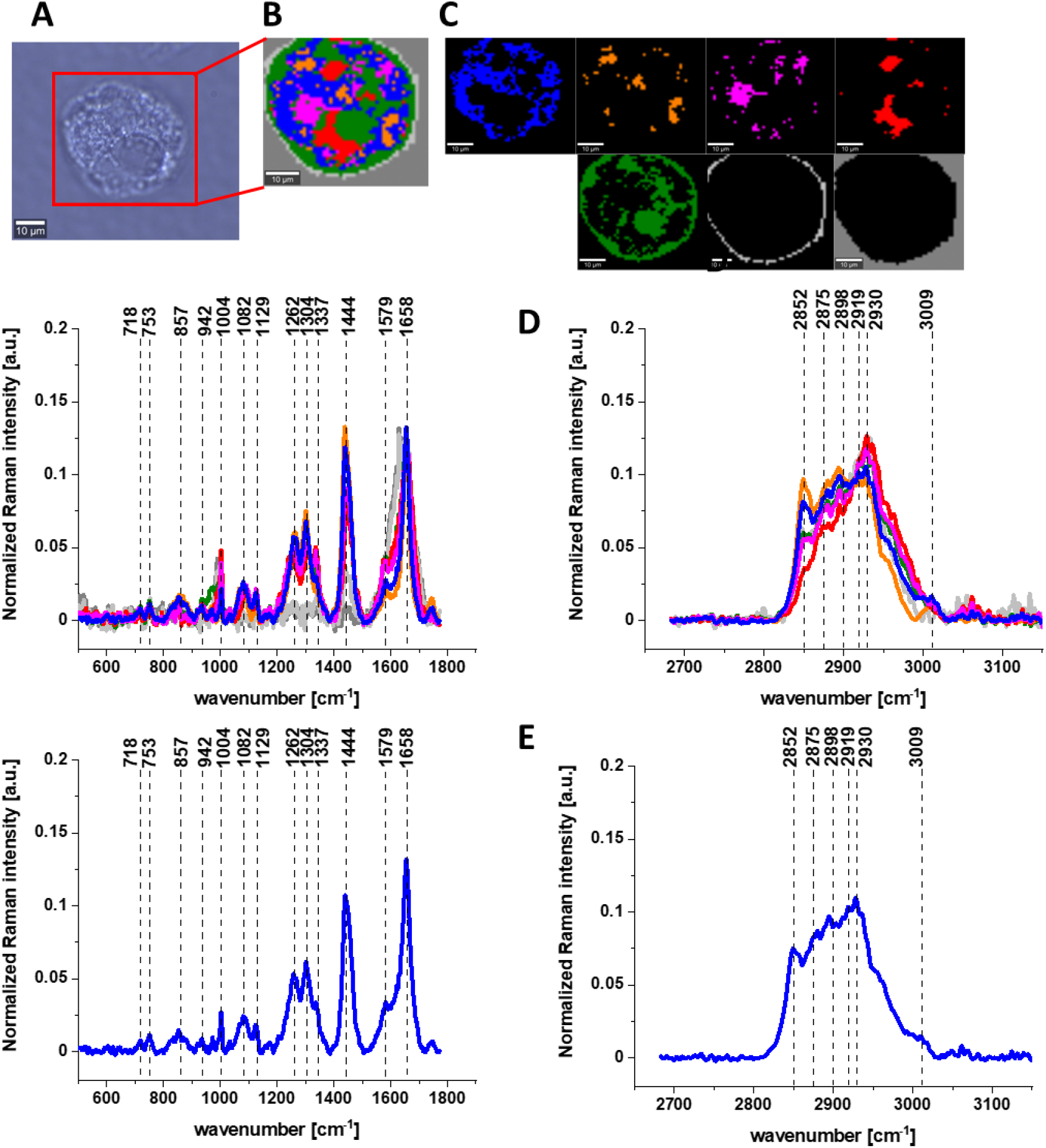
The microscopy image (A), Raman image (B) constructed based on Cluster Analysis (CA) method, Raman images of all clusters identified by CA assigned to: lipid-rich regions (blue and orange), mitochondria (magenta), nucleus (red), cytoplasm (green), cell membrane (light grey), and cell environment (dark grey) (C), the average Raman spectra typical for all identified clusters for low frequency and high frequency region (D), and the average Raman spectrum for cells as a whole (E) for cancer human colon cells after adding of mevastatin in 10 μM concentration for 48h. Cell measured in PBS, colors of the spectra correspond to the colors of clusters, excitation laser line 532 nm, the average Raman spectra calculated based on the data for 6 cells.

To check the influence on the concentration of the mevastatin the same analysis was performed for CaCo-2 cancer human colon cell line incubated with mevastatin in 50 μM concentration for 24h. Figure 4 shows the microscopy image, Raman image, Raman images of all cell substructures identified by using Cluster analysis algorithm, average Raman spectra typical for identified: lipid rich structures, mitochondria, nucleus, cytoplasm and cell membrane, cell environmental, the average Raman spectra for the cell as a whole of cancer human colon cells after adding of mevastatin in the concentration of 50 μM for 24 h.

**Figure 4.**
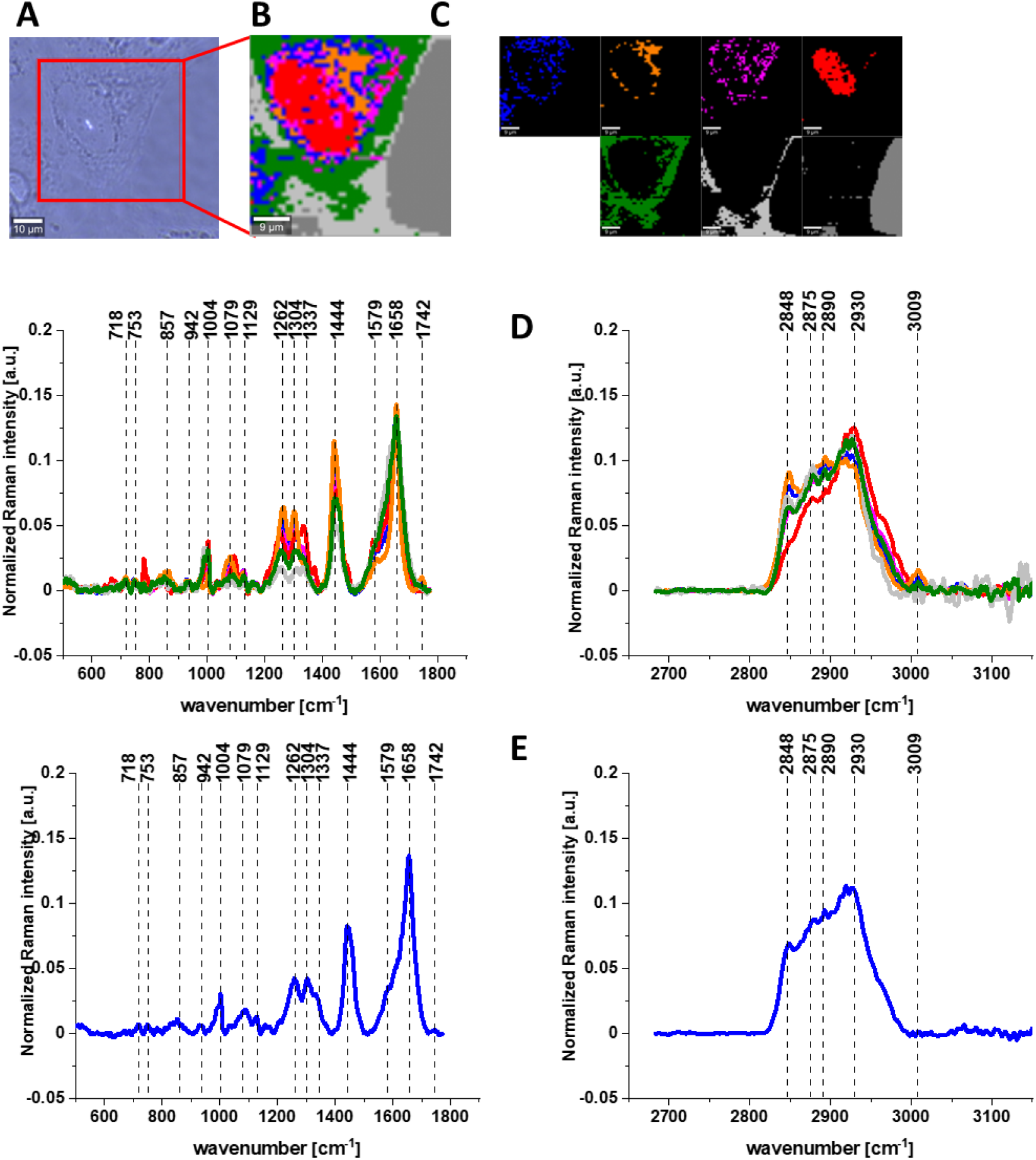
The microscopy image (A), Raman image (B) constructed based on Cluster Analysis (CA) method, Raman images of all clusters identified by CA assigned to: lipid-rich regions (blue and orange), mitochondria (magenta), nucleus (red), cytoplasm (green), cell membrane (light grey), and cell environment (dark grey) (C), the average Raman spectra typical for all identified clusters for low frequency and high frequency region (D), and the average Raman spectrum for cells as a whole (E) for cancer human colon cells after adding of mevastatin in 50 μM concentration for 24h. Cell measured in PBS, colors of the spectra correspond to the colors of clusters, excitation laser line 532 nm, the average Raman spectra calculated based on the data for 3 cells.

Based on the Raman data obtained for normal, cancer and supplemented by mevastatin cancer cells we can compare the vibrational futures of human colon cells using the average spectra calculated for cells as a whole and Raman band intensities ratios calculated for the main building blocks of biological samples: proteins, nucleic acids and lipids.

Figure 5 shows the comparison for Raman average spectra for normal CCD-18Co and cancer CaCo-2 cells of human colon.

**Figure 5.**
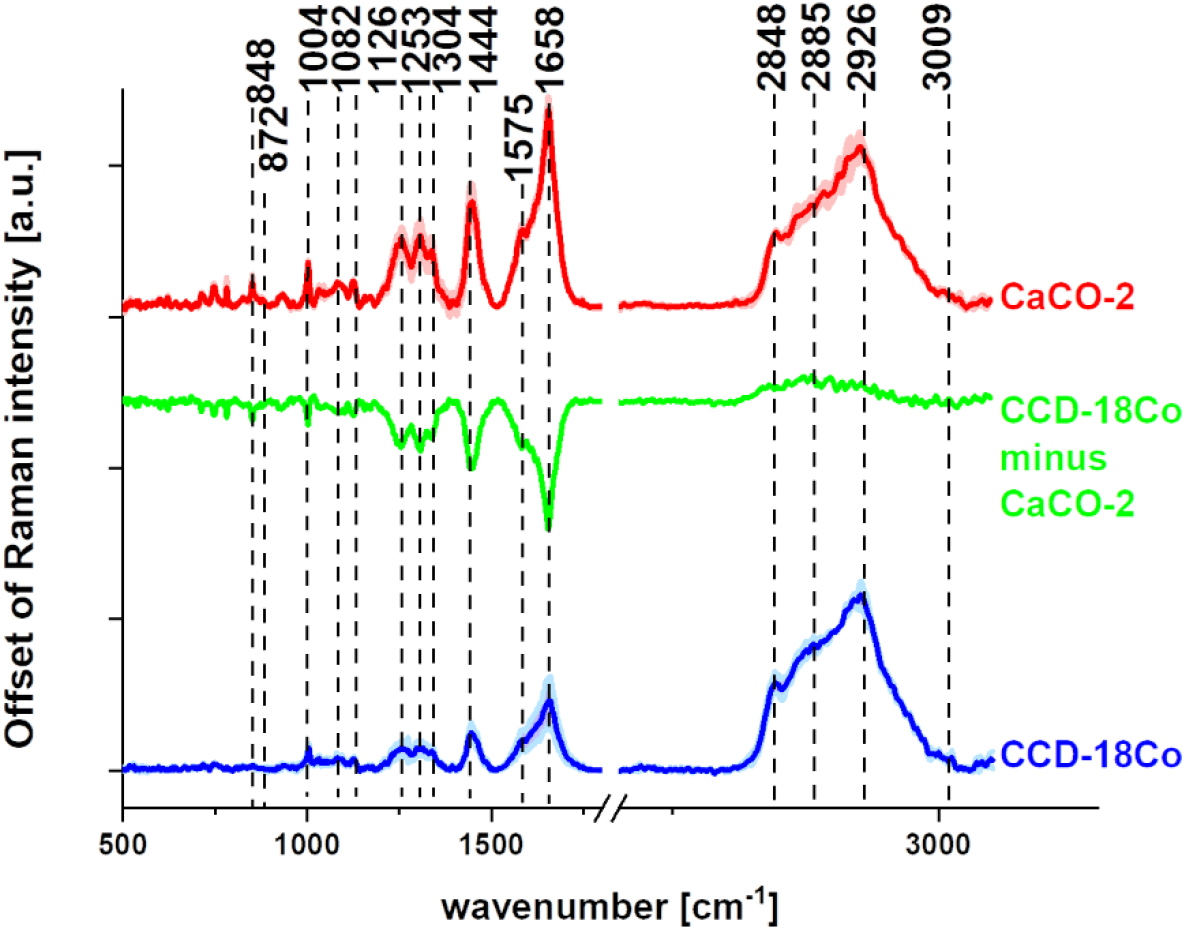
The average spectra +/− SD (SD - standard deviation) obtained for normal CCD-18Co human colon cells (blue line), the average spectra +/− SD obtained for cancer CaCo-2 human colon cells (red line) and the differential spectrum calculated based on them (green line).

For cells not supplemented by mevastatin the main differences between normal and cancer cells can be observed for the Amide III band (1230-1300cm^−1^) an the Amide I band (1600-1690 cm^−1^), whose are widely used to study the secondary structure of proteins and the global amount of proteins [45–48]. For other proteins related bands observed at 1004, 1585 and 2926 cm^−1^ also significant differences can be noticed. For CaCo-2 cells the higher content of RNA/DNA was also confirm. An increasing number of studies demonstrate e.g. the potential use of cell-free DNA as a surrogate marker for multiple indications in cancer, including diagnosis, prognosis, and monitoring of cancer [49]. The same trend can be observed for the bands 848 cm^−1^ and 850 cm^−1^ assigned to the mono- and disaccharides and tyrosine respectively [38,50]. Effect related to saccharides could be explained by the higher concentrations of glycolysis intermediates such as acetate and lactate and an increased secretion of hyaluronan [50,51]. Bands associated with the phosphates c.a. 753, 872, 1082, 1575 cm^−1^ also show a higher contribution for cancer CaCo-2 cell line. The higher phosphorylation status of cancerous tissues has been shown in the literature for many organs including breast, brain and colon [38]. In contrast to positive correlation for proteins and phosphorylated proteins observed for cancer cells the negative correlation is observed for lipids peaks from the high frequency region (bands from the region 2845-3009 cm^−1^). This finding confirm the reprograming lipids metabolism CaCo-2 cancer cells [48,52,53].

The next step in the analysis was the comparison of vibrational features obtained for not supplemented human colon cells - normal and cancer and cancer cells incubated with mevastatin for 24 or 48h shown in Figures 2–4. The obtained results are presented in Figure 6 for proteins, nucleic acids and lipid respectively.

**Figure 6.**
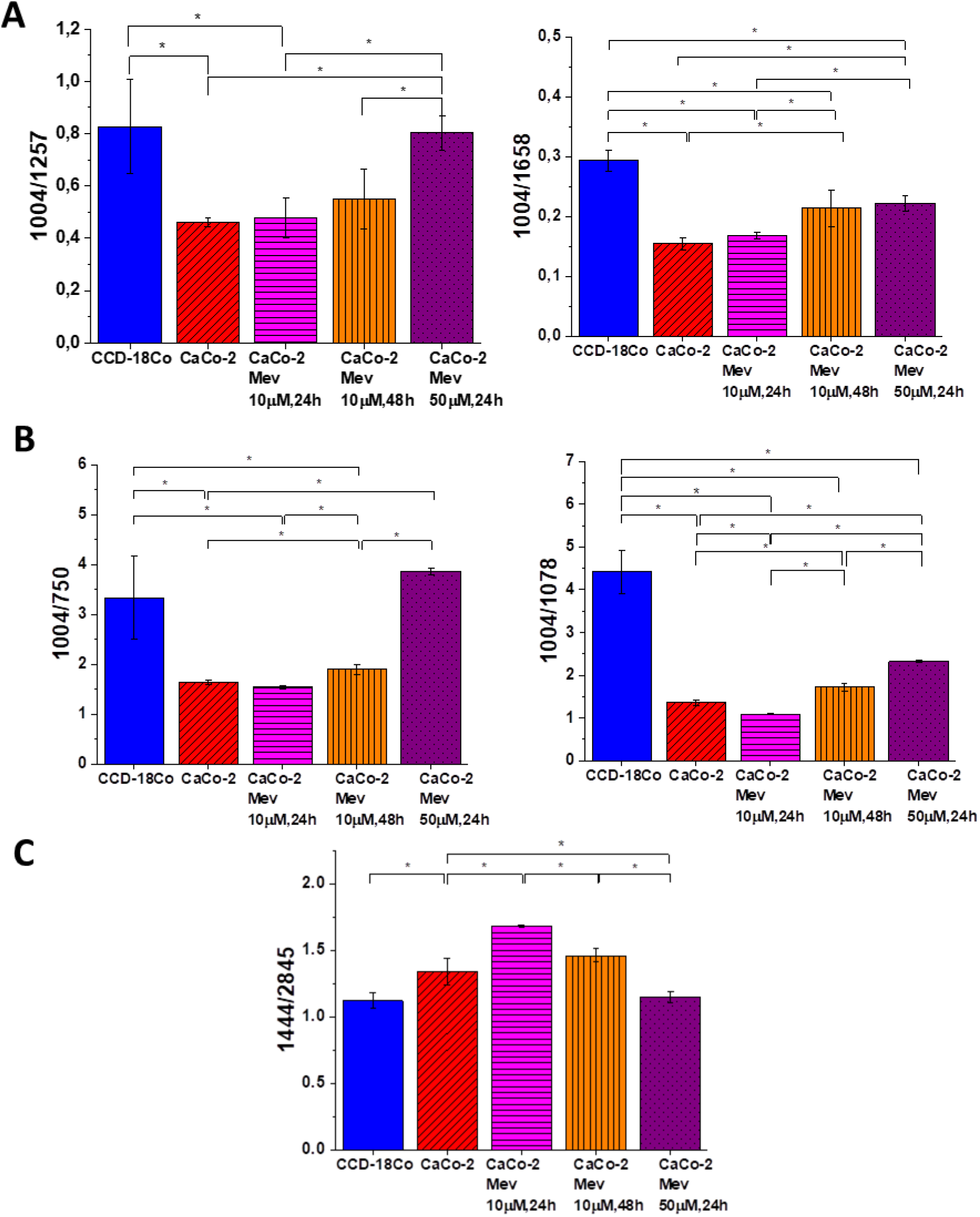
Raman band intensities ratios for selected Raman bands corresponding to proteins 1004/1257, 1004/1658 for five groups of human colon cells: normal human colon cells CCD18-Co - control group (labeled CCD-18Co, blue), cancer human colon cells CaCo-2 (labeled with CaCo-2, red), cancer human colon cells CaCo-2 incubated with mevastatin in concentration of 10 μM for 24 h (labeled with CaCo-2 Mev 10mM,24h, magenta), cancer human colon cells CaCo-2 incubated with mevastatin in concentration of 10 μM for for 48 h (labeled with CaCo-2 Mev 10μM,48, orange) and cancer human colon cells CaCo-2 incubated with mevastatin in concentration of 50μM for for 24 h (labeled with CaCo-2 Mev 50μM,24h, violet), statistically significant data, based on ANOVA analysis, were marked with asterix (*).

Based on the data obtained by using Raman spectroscopy and imaging for proteins one can see that synthesis of this class of biological building compounds was modulated by the adding of mevastatin and was also differentiating between normal and cancer cells not supplemented by mevastatin. For CaCo-2 cancer cells we observed the lower intensities for ratios 1004/1257 and 1004/1658. Such results were expected taking into account that the development of cancer is associated with the overexpression of proteins. However, for cancer cells incubated with mevastatin in 10μM concentration we noticed the statistically significant increase of analyzed ratios, which suggest that statin-induced inhibition of protein synthesis can be an underlying mechanism for statin-induced cell death [54]. Protein synthesis is one of the most complicated biochemical processes undertaken by the cell, requiring approximately 150 different polypeptides and 70 different RNAs. In addition protein synthesis can be halted when only a small fraction of the ribosome is inactivated by certain ribotoxins [55]. Alternatively activation of various stress related kinases within the cell could reduce the rates of protein synthesis [56]. Some pathways are more specific to the actions of statins. HMG-CoA reductase is associated with the endoplasmic reticulum (ER) [57], which regulates protein synthesis and ER stress leads to apoptosis [58,59].

Based on the data presented in Figure 6 for bands typical for nucleic acids one can see that the intensity of the ratios typical for these compounds 1004/750 and 1004/1078 decreases for CaCo-2 human cancer colon cells compared to control group - CCD-18Co corresponding to the normal human colon cells. This observation confirm that the synthesis of nucleic acids in cancer cells is enhanced compared to normal cells, which is the expected result. Moreover, analyzing Figure 6 one can noticed that the adding of mevastatin to the culture medium affects the amount of nucleic acids observed in CaCo-2 cancer cells and the ratios typical for lower concentration and longer time of incubation (labeled with CaCo-2 Mev 10μM,48, orange) or higher concentration and shorter time of incubation (labeled with CaCo-2 Mev 50μM,24h, violet) inhibits the synthesis of nucleic acids and the same reduces the proliferation ability of CaCo-2 cells. This finding is supported by scientific literature confirming, using traditional molecular biology methods, that decreased levels of DNA for cells interacting with statins is observed [19,60–62].

The statistically significant differences between normal human colon cells, cancer human colon cells and cancer human colon cells supplemented by using mevastatin have been found also for lipids components of analyzed samples. It is well known from the literature that statins modulate the lipids composition of cells and tissues due to the influence on cholesterol level (in general, statins represent HMG-CoA reductase inhibitors, and are widely used for the treatment of hypercholesterolaemia) and the reduction of triglyceride concentrations. Results obtained based on intensity of Raman peaks related to lipids (1444, 2854, 3009 cm^−1^) confirmed that decreasing of intensity of peaks typical for lipids for cells treated by mevastatin is observed and this effect is time and dose dependent [63].

Investigations regarding cells biochemistry were extended with analysis of nanomechanical properties of human colon cells: normal, cancer and cancer supplemented by mevastatin by Atomic Force Microscopy. Figure 7 presents the data obtained during AFM measurements: topography maps, curves related to the topography measurements for forward and backward traces (for randomly chosen point of cell), topography maps with marked area for which the force-distance curves were measured and analysis of Young’s modulus, adhesion force and intendation calculated based on force-distance curves for all analyzed types of samples.

**Figure 7.**
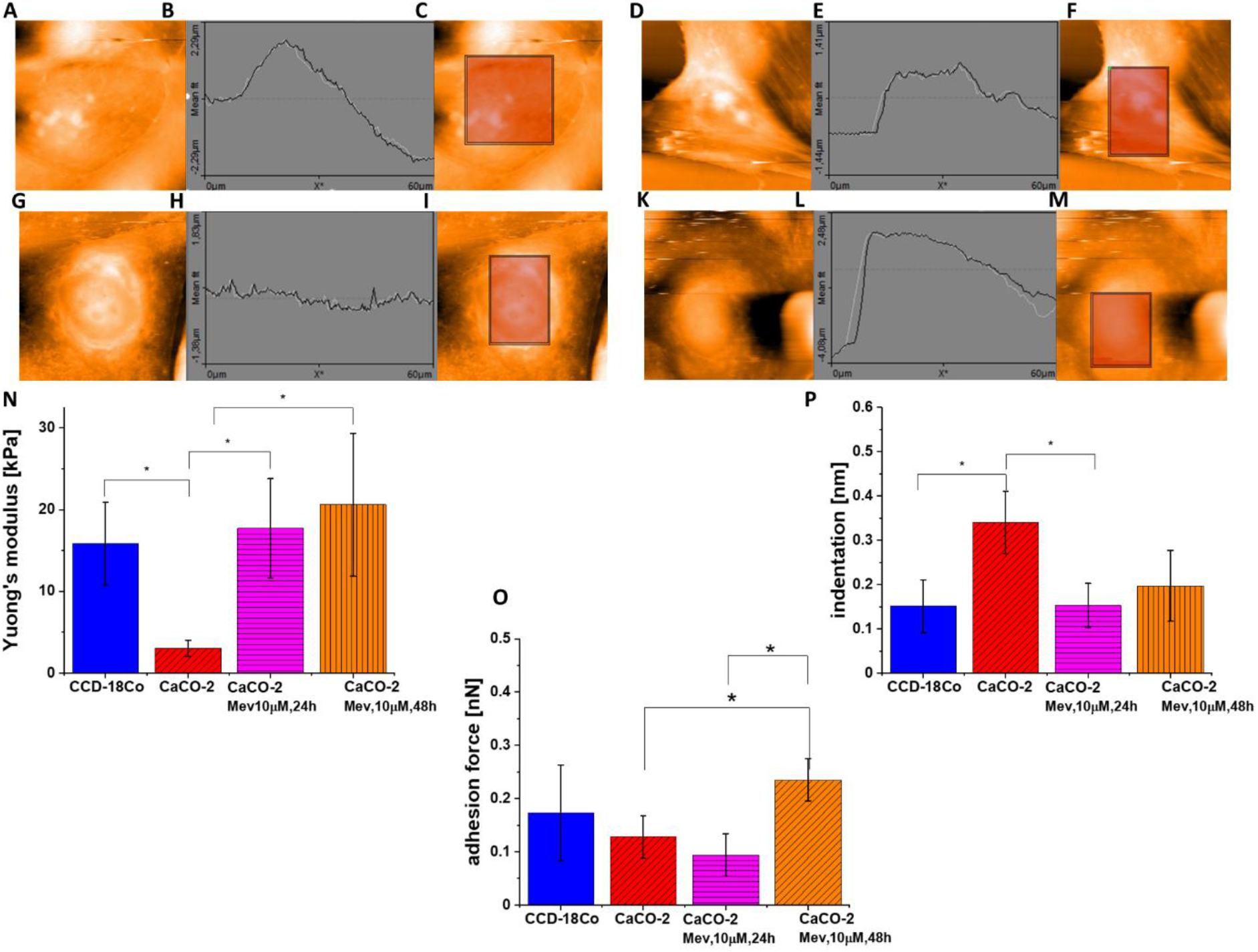
AFM topography maps of CaCo-2 (A), CCD-18Co (D), CaCo-2 supplemented by mevastatin 10 μM, 24h (G), and CaCo-2 supplemented by mevastatin 10 μM, 48h (K), curves related to the topography measurements for forward and backward trace of CaCo-2 (B), CCD-18Co (E), CaCo-2 supplemented by mevastatin 10 μM, 24h (H), and CaCo-2 supplemented by mevastatin 10 μM, 48h (L), topography maps with marked area for which the force-distance curves were measured of CaCo-2 (C), CCD-18Co (F), CaCo-2 supplemented by mevastatin 10 μM, 24h (I), and CaCo-2 supplemented by mevastatin 10 μM, 48h (M) and Young’s modulus (N), adhesion force (O) and indentation (P) calculated based on force-distance curves for all analyzed types of samples.

One can see from Figure 7 that the cancer human colon cells CaCo-2 are more elastic compared to normal human colon cells CCD-18 Co and the adding of mevastatin in 10 μM concentration which is time depending changes the nanomechanical properties of these cells. The supplementation by using mevastatin brings the cancer cell closer in elasticity to normal CCD-18 human colon cells, which confirm the changes in skeleton organization of analyzed cells. The obtained result is consistent with literature data, which confirm higher flexibility of cancer cells compared to normal one [64].

## Conclusions

The studies performed with the use of Raman spectroscopy and imaging allowed to characterize the biochemical composition of normal CCD-18Co and cancerous CaCo-2 cells in human colon. Interpretation of the results carried out using the Cluster Analysis method made it possible to obtain average Raman spectra, both for cells as a single cluster and for individual cell organelles. Differences between CCD-18Co and CaCo-2 cells, based on the analysis of Raman spectra, were identified for bands typical of nucleic acids, proteins and lipids. The addition of mevaststin for 24h and 48h modulated the metabolism of Caco-2 cancer cells in the human colon, and the recorded Raman spectra confirmed the changes of intensity on bands typical for proteins, nucleic acids and lipids. The changes for main cells building compounds for mevastatin supplementation cells have shown also the dose effect of statin adding.

Research carried out using Atomic Force Microscopy - AFM allowed to characterize the nanomechanical properties of normal CCD-18Co cells and cancerous CaCo-2 cells of the human colon without supplementation and after adding mevaststin for a period of 24 and 48 hours. Based on the investigation performed, it can be concluded that cancerous cells are characterized by a lower value of Young’s modulus than normal cells, respectively 3.04±0.98 kPa and 15.84±5.06 kPa. The addition of mevaststin caused a increase in the value of Young’s modulus by about 80 %, which confirms the influence of mevaststin on the organization of the cellular cytoskeleton.

The use of Atomic Force Microscopy to characterize elastic properties of normal and cancer cell line justifies the idea of using this method to track the changes of morphological and topographical appearance with the progression or regression of the tumor.

## Author Contributions

Conceptualization: BB-P; Funding acquisition: BB-P; Investigation: KB, BB-P; Methodology: BB-P, KB, Writing - original draft: KB, BB-P; Manuscript editing: KB, BB-P. All authors reviewed and provide feedback on the manuscripts. All authors have read and agreed to the published version of the manuscript.

## Acknowledgments

This article has been completed while the first author was the Doctoral Candidate in the Interdisciplinary Doctoral School at the Lodz University of Technology, Poland.

